# Quantitative clonal analysis reveals vast heterogeneity among fallopian tube cells for ovarian cancer-initiation and progression

**DOI:** 10.1101/2022.01.23.477434

**Authors:** Jianhao Zeng, Astrid Catalina Alvarez-Yela, Eli Casarez, Ying Jiang, Lixin Wang, Brianna E. Kelly, Eugene Ke, Taylor Jenkins, Kristen A. Atkins, Kevin A. Janes, Jill K. Slack-Davis, Hui Zong

## Abstract

Different cellular compartments within a tissue present distinct cancer-initiating capacities. Current approaches to delineate cancer-initiating heterogeneity require a well-understood lineage hierarchy and cell-specific genetic tools, which are lacking for many tissues. Here, we circumvented this hurdle and investigated the capacity of fallopian tube cells in initiating serous ovarian cancer, utilizing a mouse genetic system that generates scattered GFP-labeled mutant cells. Through quantitative tracing of clonal expansion of individual mutant cells, we revealed that only a rare primitive subset enriched in the distal fallopian tube is capable of clonal expansion whereas others stall immediately upon acquiring oncogenic mutations. Expanded clones from this subset then encountered further attrition: many stalled shortly after while others sustained small cluster of proliferative cells and biased differentiation toward primitive fate. Taken together, our study showcases quantitative clonal tracing as a powerful approach to investigate cancer-initiating heterogeneity and clonal trajectory in tissues with limited prior knowledge.

## Introduction

Cancer initiation is a fundamental question in cancer biology. While oncogenic mutations are key for cancer initiation, a permissive context from the cells where these mutations occur is also indispensable. Different cellular compartments within a tissue could manifest vastly different cancer-initiating capacities even if hit by the same oncogenic events (Perez-Losada and Balmain, 2003). For example, APC deletion in mouse intestinal stem cells initiates cancer and causes adenomas in 3-5 weeks. However, when the same mutation occurs in the transit-amplifying cells, adenomas fail to form, suggesting that intrinsic properties of stem cells uniquely permit cancer initiation upon APC loss (Barker et al., 2009). The tumorigenic potential of distinct cell types can be rigorously interrogated with mouse models when precise conditional genetic targeting tools for each cell type are available. However, this approach is often hampered by our limited knowledge of tissue lineage hierarchy and the lack of cell-specific Cre transgenes.

To dissect cellular heterogeneity of cancer-initiating capacity in tissues lacking an established lineage hierarchy, we deploy a mouse genetic system called Mosaic Analysis with Double Markers (MADM). MADM induces scattered single mutant cells in a broad category of cells under the control of a pan-tissue Cre transgene and simultaneously labels these cells with GFP. Via quantitative tracing of the clonal expansion of individual mutant cells, we can pinpoint those highly expanded clones originating from cells with cancer-initiating capacity whose molecular features can be further interrogated through spatial profiling. MADM contains a pair of knock-in cassettes of chimeric GFP and RFP coding sequences separated by LoxP sites. These cassettes are associated with the wildtype or mutant alleles of tumor suppressor genes on the same chromosome. Through Cre/loxP-mediated inter-chromosomal mitotic recombination that occurs at a low frequency (0.1%–1% or even lower), MADM generates sporadic mutant cells labeled with GFP (**Fig. 1*A_1_***) (Muzumdar et al., 2007; Zong et al., 2005). The rarity of mutant cells and the unequivocal GFP-labeling of mutated cells provide unprecedented clonal resolution to trace the expansion of individual mutant cells (**Fig. 1*A_2_***) (Hippenmeyer, 2013; Liu et al., 2011; Zong et al., 2005).

**Figure 1.**
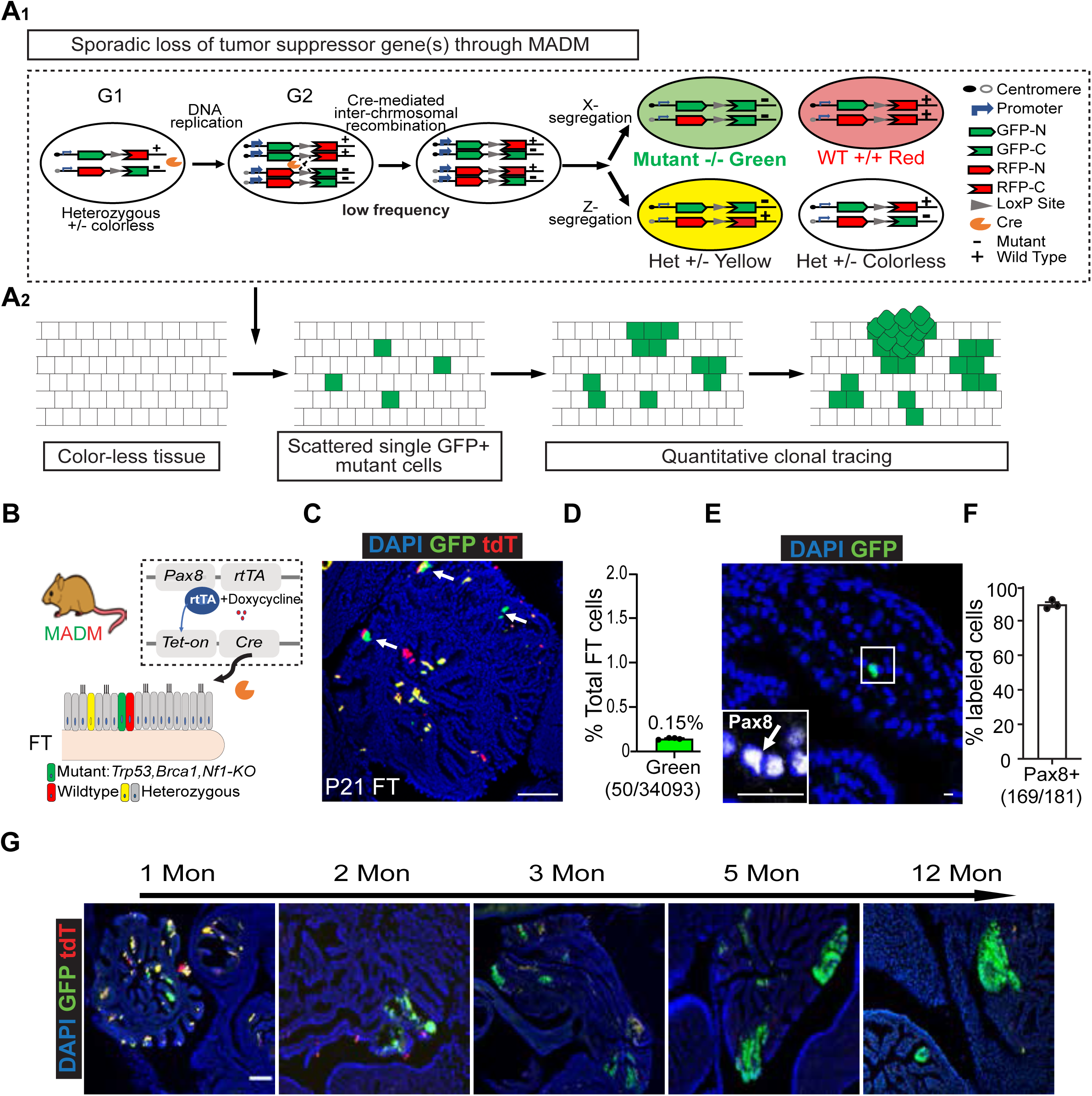
Genetic marking of single mutant cells at clonal density through MADM in FT Pax8+ Cells. (A) A_1:_ MADM mechanism: MADM contains a pair of knock-in cassettes of chimeric GFP and RFP (tdTomato, tdT) coding sequences, separated by a loxP-containing intron. Cre-mediated inter-chromosomal recombination in mitotic cells followed by X segregation generates a single GFP+ cell that is a homozygous mutant, and a sibling tdTomato+ cell that is wildtype; or by Z segregation which generates a GFP+ & tdTomato+ double positive heterozygous cell and a sibling colorless heterozygous cell. A_2:_ MADM could be used to induce sporadic single GFP+ mutant cells in the tissue, which allows quantitative tracing of mutant cell progression at the clonal resolution. (B) The scheme of a MADM-based mouse model with scattered single GFP+ mutant cells generated in the FT. (C) MADM-labeled cells in the FT after inducing Cre between postnatal day 0 and day 21 (P0-21). Scale bar=100µm. (D) The frequency of GFP+ cells in the FT after inducing Cre activity (±SEM, *n*=4). (E) MADM-labeled cells were stained positive for Pax8. Scale bar=50µm. (F) Pax8+% among all MADM-labeled cells. (*n* =3). (G) Expansion of GFP+ mutant clones over time.

Utilizing this research paradigm, we sought to dissect the cancer-initiating capability of the fallopian tube (FT) Pax8 cells, which have emerged as a cell of origin for high-grade serous ovarian cancer (HGSOC). HGSOC is the most prevalent and aggressive ovarian cancer subtype, which causes about 14,000 deaths annually in the United States (Siegel et al., 2020; Vaughan et al., 2011). While HGSOC was initially thought to arise solely from the ovarian surface epithelium, an increasing body of epidemiologic, clinical, molecular, and mouse studies suggest that the fallopian tube (FT) is an alternative origin for a considerable portion of HGSOC. In human studies, FT lesions (serous tubal intraepithelial carcinomas, STICs) that resemble ovarian tumors were frequently detected (Kindelberger et al., 2007; Lee et al., 2007; Medeiros et al., 2006; Przybycin et al., 2010; Visvanathan et al., 2018). These FT lesions share identical *TP53* mutations (Ardighieri et al., 2016; Kuhn et al., 2012; Labidi-Galy et al., 2017; Soong et al., 2018) and similar transcriptomic profiles with concurrent HGSOC tumors (Beirne et al., 2019; Ducie et al., 2017; Marquez et al., 2005; Tone et al., 2008), indicating their close lineage relationship. Moreover, surgical removal of FTs (salpingectomy) prominently reduces ovarian cancer risk by 42-50% (Falconer et al., 2015; Madsen et al., 2015). In studies with genetically engineered mouse models, the transformation of epithelial cells in the oviduct (analogous to the fallopian tubes in humans, referred to as fallopian tube hereafter) with clinically relevant mutations (*TP53, BRCA1, RB1, PTEN, NF1, etc.*) directly leads to FT lesions and ovarian tumors that resemble human disease (Perets et al., 2013; Sherman-Baust et al., 2014; Wu et al., 2016; Zhang et al., 2019). More and more evidence is accumulating in support of the fallopian tube as an origin of HGSOC.

The FT is known to consist of ciliated cells (Acetyl-Tubulin+) that facilitate gamete transportation and secretory cells (Pax8+) that produce nutrient-rich fluid (Stewart and Behringer, 2012). The latter (Pax8+) serves as the cancer cell-of-origin as they constitute most HGSOC tumor cells. Moreover, the direct transformation of Pax8+ cells but not ciliated cells in transgenic mice leads to HGSOC (Karnezis et al., 2017; Perets et al., 2013). However, recent studies suggest heterogeneity among FT Pax8+ cells. As in vitro cultures of isolated FT epithelial cells, Pax8+ cells have a heterogeneous ability to form organoids and self-renew, indicating the existence of a more primitive subset (Paik et al., 2012; Rose et al., 2020). Although in vivo data are lacking, these findings raise the question of whether all or only specific subsets of Pax8+ cells possess the cancer-initiating capacity in vivo. Addressing this question should further our understanding of HGSOC initiation and lay the foundation for earlier detection and prevention. However, such efforts have been hindered by the lack of highly specific markers that can discriminate the subset with high organoid-forming capability from those with low capability.

Here, taking advantage of MADM’s strengths in generating scattered single mutant cells to enable quantitative clonal tracing, we interrogated the cellular heterogeneity of FT Pax8+ cells in their capacity to initiate cancer. We found that only a rare primitive subpopulation enriched in the distal FT provides the permissive context for cancer initiation. We also delineated the early progressive trajectory from initiation toward FT lesions. Therefore, quantitative clonal tracing with MADM presents as an excellent solution for assessing heterogeneity in cancer-initiating potency in tissues lacking well-defined lineages.

## Results

### Genetic marking of single mutant cells at clonal density through MADM in FT Pax8+ population

To generate and track individual mutant cells within the FT Pax8+ population through MADM, two critical prerequisites need to be considered: 1) a Cre transgene that specifically targets the FT Pax8+ population, 2) HGSOC-relevant tumor suppressor genes reside on the telomeric side of the MADM cassettes. We incorporated the *Pax8-rtTA; TetO-Cre* system to target the FT Pax8+ population under the control of doxycycline (Dox) (Perets et al., 2013; Traykova-Brauch et al., 2008). To select the clinical-relevant tumor suppressor genes, we analyzed TCGA datasets of human HGSOC patients. We found that the loss of *TP53* (63%), *BRCA1/2* (78%), and the mutations that activate the Ras-MAPK pathway (37%, partially through loss of *NF1*) are among the most prevalent genetic alterations and often co-occur (**Fig. S1*A***) (Ahmed et al., 2010; Network, 2011; Patch et al., 2015; Toss et al., 2015; Walsh et al., 2011). Since *Trp53, Brca1,* and *Nf1* in mice all reside on chromosome-11, where the MADM cassettes had been knocked in (**Fig. S1*B***) (Hippenmeyer et al., 2010; Tasic et al., 2012) we selected these mutations to drive tumor initiation. After multi-generational breeding to incorporate the Cre transgene and the mutations into two MADM stock lines (**Fig. S1*C***, see methods for details), we generated MADM-mutant mice in which single GFP+ mutant cells were generated within the FT Pax8+ population (**Fig. 1*B***). This breeding scheme also generated MADM-wildtype mice for single-cell clonal tracing of wildtype cells (**Fig. S1*C***).

To assess whether the frequency of GFP-marked cells is low enough for clonal tracing, we induced Cre activity between postnatal day 1 and 21 (P0-21) with Dox in the drinking water and then assessed GFP+% among all FT epithelial cells. We found that less than 0.2% of FT epithelial cells were GFP+, a sparseness that allows clonal tracing (**Fig. 1*C, D***). RFP+ wildtype cells were generated at a similar frequency, whereas double-positive (yellow) heterozygous cells were generated at ~8-fold higher rate due to G2-Z segregation during mitosis plus G0/G1 recombination (**Fig. S2*A, B***). We will mainly focus on analyzing the GFP+ mutant cells hereafter. We also found nearly all MADM-labeled cells were Pax8+ (**Fig. 1*E, F***), which verified the faithful expression of Pax8-rtTA; TetO-Cre transgenes. As mice aged, the GFP+ mutant cells gradually formed clones that occupy a continuous stretch of FT epithelium, indicating the capability of the initial single mutant cells to undergo clonal progression (**Fig. 1*G***). Some clones eventually formed FT STICs lesions featured by foci of atypical columnar epithelial cells with enlarged nuclei, loss of cilia, and increased proliferation (**Fig. S2*C, D***). Although aging animals beyond one year to assess the onset of full-blown HGSOC was not possible because mice succumbed to invasive tumors in the uterus, which also contains Pax8+ cells with transforming potential (Tacha et al., 2011) (**Fig. S2*E, F***), this model is uniquely suitable for clonal tracing of individual mutant FT Pax8+ cells to dissect their cancer-initiating capacity.

### Dichotomous expansion of individual mutant cells revealed by quantitative clonal tracing

Next, to determine whether all Pax8+ cells share similar cancer-initiating capacity, we evaluated the premalignant clonal expansion of individual mutant Pax8+ cells before STIC formation. We first induced Cre activity between P0-21 to generate GFP+ mutant cells at clonal density and then analyzed their clonal expansion at five months of age (**Fig. 2*A***), when initiated cells have expanded but not yet formed histopathologically detectable lesions. To gain a complete view of all clones, we optically cleared the FTs with the CUBIC method (Susaki et al., 2015), imaged the entire FT with light-sheet microscopy, and then quantified the size of each clone by reconstructing image stacks (**Fig. 2*B***). Interestingly, clone sizes were highly divergent: ~75% of the clones contained < 10 cells, yet a small fraction of clones expanded to hundreds of cells or more (**Fig. 2C, D**). To exclude the possibility that the dichotomy resulted from stochastic fluctuations, we implemented a mathematical analysis that evaluates whether the observed clonal size distribution can be explained by one single stochastic group. We estimated the clonal size distribution from the majority clones and asked whether the frequency of observed outlier large clones is within that predicted by the majority distribution. The observed clonal sizes were best fit by a negative binomial model for clones below 50 cells (KS test, p-value<0.1, see methods for details) (Bliss and Fisher, 1953). Under the null hypothesis, this one-group model predicts one clone larger than 50 cells. In contrast, we observed 27 clones larger than 50 cells, suggesting the existence of an outlier group (**Fig. 2*E***) (*p* < 10^−8^, Fisher Exact Test) that has a higher expansion potential than most clones.

**Figure 2:**
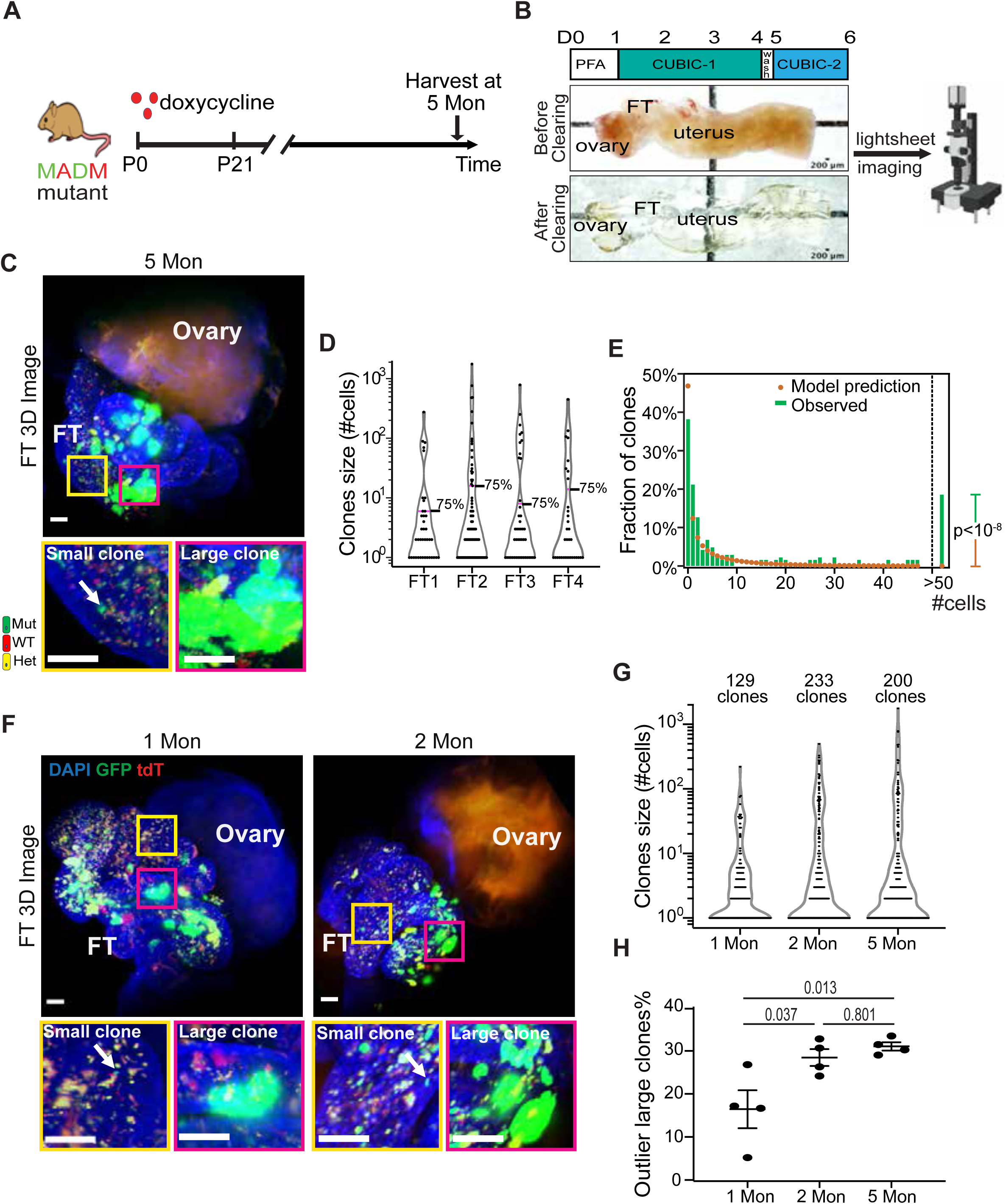
Dichotomous expansion of individual mutant cells revealed by quantitative clonal tracing. (A) The scheme of DOX treatment and FT harvesting for measuring the size of mutant clones. (B) The pipeline for high-resolution whole-FT imaging: fixed reproductive tracts were cleared through the CUBIC method, followed by 3D imaging with light-sheet microscopy. (C) Representative 3D images of cleared whole FT from 5 months old MADM-mutant mice (*n*=4). The yellow box shows non-expanded clones, the magenta box shows expanded clones. Scale bar=50 µm. (D) Violin plot of the clonal size distribution of mutant clones in four independent FTs from mice at 5 months of age. (E) Fitting the observed clonal size distribution of mutant clones at 5 months to a negative binomial model. Bars show observed data, points show model prediction. The number of observed clones >50 cells is significantly higher than predicted. Fisher Exact Test, p < 10^−8^. (F) Representative 3D images of cleared whole FT from 1 and 2 months old MADM-mutant mice (*n*=4). The yellow box shows non-expanded clones, the magenta box shows expanded clones. Scale bar=50 µm. (G) Violin plot of the clonal size distribution of mutant clones at 1 and 2 months. Data pooled from 4 mice. (H) The portion of profoundly-expanded outlier clones at 1,2,5 months (± SEM, *n* = 4, ANOVA test with Tukey HSD).

To determine when these outlier clones become detectable, we repeated the clonal size measurements with mice at one and two months of age. To our surprise, profoundly expanded clones already existed among minute ones shortly after Cre induction (**Fig. 2*F, G***). We ruled out the trivial explanation that this clonal size divergence simply reflects their differential birth date over the 21 days of doxycycline administration (P0-21) by demonstrating prominent clonal size variance even when the clonal birth date was limited within a 3-day time window (**Fig. S3*A, B, C***). To further check whether the clonal size dichotomy merely reflects fluctuation of the hormone at different estrus cycles (Rendi et al., 2012), we harvested post-puberty mice at the proestrus phase (peak of estrogen, valley of progesterone) or diestrus phase (valley of estrogen, peak of progesterone). We found that, in both phases, the clonal size of mutant clones showed prominent dichotomy and the average size of highly expanded clones was comparable (**Fig. S3*D, E***). These data indicate that dichotomous clonal expansion of mutant Pax8+ cells is not due to the short-term fluctuation of steroid hormones during the estrous cycle. Finally, we compared the frequency of expanded large clones from 1, 2, and 5 months of age. Because the variance of clonal sizes at younger ages was smaller overall, a Poisson model was used to calculate the portion of outlier large clones (see methods for details). We found that the portion of outlier large clones at one month is 16.5 ± 4.4%, which increased to 28.4 ± 1.9% at two months and held steady at 30.9 ± 0.9% at five months (**Fig. 2*H***), suggesting that the expansion potential of initiated Pax8+ cells was determined early. Collectively, these data show that initiated FT Pax8+ cells present dichotomous expansion potential with the majority immediately stalled and a small fraction profoundly expanded.

**Figure 3:**
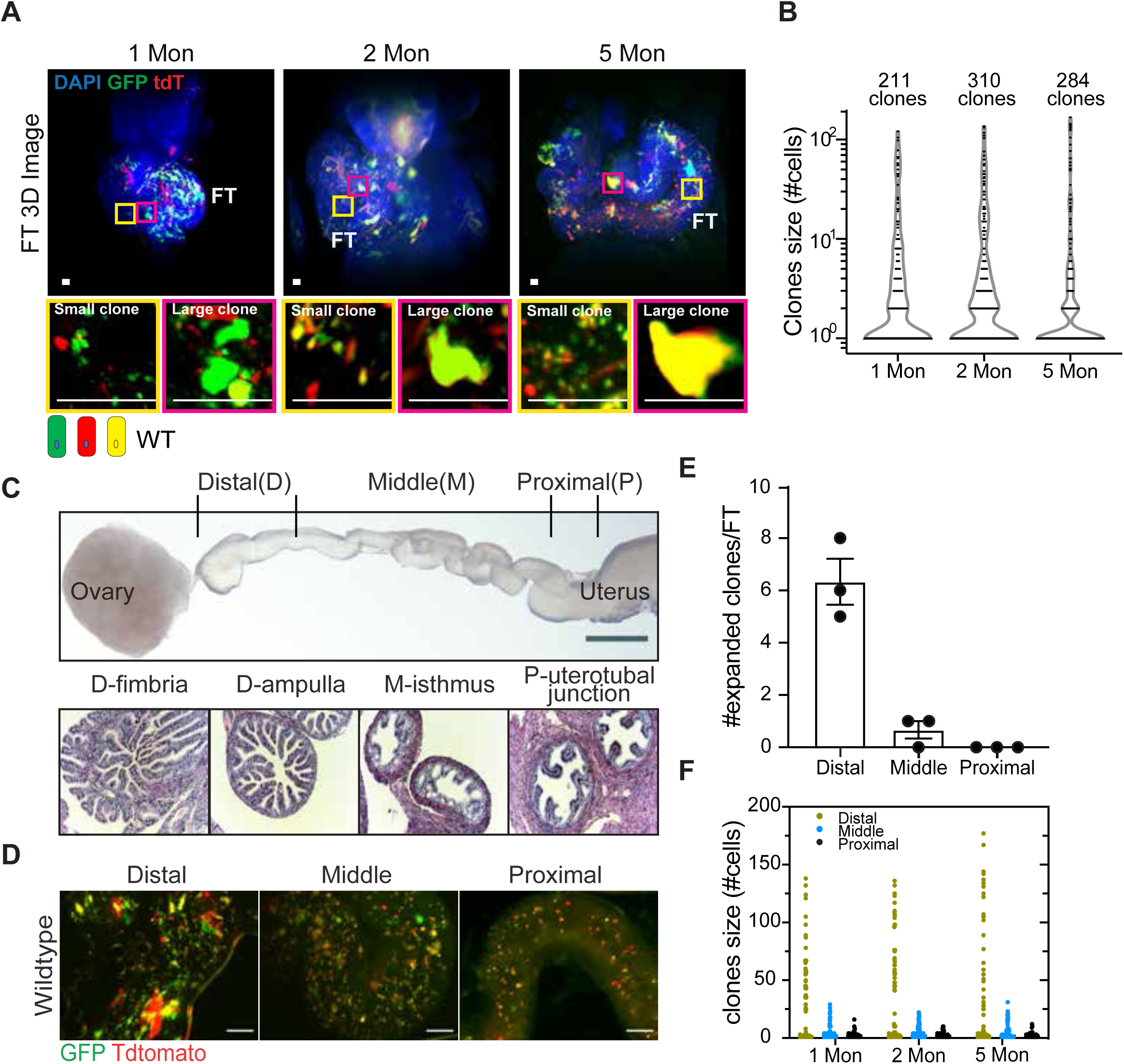
Dichotomous clonal expansion occurred among the Pax8+ population even in the absence of oncogenic mutations. (A) Representative 3D images of cleared whole FT from 1,2,5 months old MADM-wildtype mice (n=3). The yellow box shows non-expanded small clones, the magenta box shows expanded large clones. Scale bar=25µm. (B) Violin plot of the clonal size distribution of wildtype clones at 1,2,5 months. Data pooled from 3 mice. (C) Upper panel: Whole-mount white-field image of a stretched FT from one-month-old MADM-wildtype mice which can be separated into three portions based on the distance to the uterus; Lower panel: each FT region has distinct tubal morphology and inner folds shape. Scale bar= 500µm. (D) Whole-mount fluorescence image of each segment of the stretched FT from MADM wildtype mice. Scale bar= 500µm. (E) The number of expanded clones in distal(D), middle(M), and proximal(P) segments of FTs from one-month-old MADM-wildtype mice. (±SEM, n=3). (F) The clonal size distribution in distal(D), middle(M) and proximal(P) segments of FTs from 1,2,5 months old MADM-wildtype mice. Data pooled from three mice at each age.

### Dichotomous clonal expansion occurred even in the absence of oncogenic mutations

The nearly immediate divergence of clonal expansion prompted us to ask whether the initial oncogenic mutations were required or if there could be intrinsic heterogeneity of the expansive potential of Pax8+ cells. To address this question, we induced clones in the MADM-wildtype mice, in which clones of all colors are free of initial oncogenic mutations. We examined the size of these wildtype clones at 1, 2, and 5 months of age and still observed evident clonal size divergence as early as 1 month of age (**Fig. 3*A, B***), indicating that the divergence of clonal expansion does not depend on the oncogenic mutations.

The FT comprises three distinct portions that can be discriminated by their distance to the ovary and their unique ductal morphology (**Fig. 3*C***). The distal FT is found to be the preferential site for cancer initiation in humans, presumably due to its proximity to the ovary, thus repeated exposure to ovulation-related inflammatory cytokines (King et al., 2011; Wu et al., 2017). We asked whether the aforementioned highly expanded large clones show similar spatial preference toward the distal end and whether their expansion is corresponding to ovulation effects. To address this question, we stretched the FTs from MADM-wildtype mice at one month and performed whole-mount imaging to examine the location of expanded clones. We found that large clones reside almost exclusively in the distal FTs, whereas stalled small clones are distributed in all three FT portions (**Fig. 3*D, E***). Similar spatial preference for large clones was also observed when we repeated the analysis with mice at two and five months old (**Fig. 3*F***), and with mutant clones (**Fig. S4*A, B, C***). Importantly, such spatial preference was not simply caused by the biased generation of more clones in the distal FT, since the initial labeling frequency was similar throughout FT segments (**Fig. S4*D***). The spatial preference of highly clonogenic cells was maintained in adult FT (Yamanouchi et al., 2010): when we induce clones between 5-6 weeks of age, large clones still mainly occurred in the distal FT regardless of their wildtype or mutant genotype (**Fig. S4*E***). The distal-FT preference of the large clones is unlikely due to ovulation, as these expanded large clones already occurred in one-month-old mice, an age when ovulation effect is limited. Instead, these data hint that a subset of FT Pax8+ cells in the distal FT may intrinsically present higher clonogenic potential that confers cancer-initiating capacity.

### Founders of highly expansive clones represent a more primitive FT population

The distal FT preference and high clonogenic potential of the founder cells for large clones raised the possibility that these cells might represent a more primitive population (Fuchs and Segre, 2000; Ghosh et al., 2017; Paik *et al*., 2012; Rose *et al*., 2020; Wang et al., 2012; Xie et al., 2018). Speculating that a portion of the cells within the large clones might retain the primitive property of the clonal founders, we hypothesized that the large clones maintain a more primitive status than the small clones. To test this hypothesis, we isolated large clones and small clones separately by fluorescence-guided laser capture microdissection. We used FTs from one-month-old MADM-wildtype mice, when large clones are readily distinguishable from small clones by size but not yet overly expanded to the extent that signatures of the primitive cells could be overshadowed by differentiated cells (Ghosh et al., 2017). From each of the six mice, we microdissected small clones and large clones to collect ~200 cells per sample. These 6 pairs of samples underwent RNA extraction, cDNA preparation, and linear amplification followed by RNA sequencing at a depth of ~ 4 million aligned reads per sample (Singh et al., 2019; Wang and Janes, 2013) (**Fig. 4*A***, see methods for details). From the RNA sequencing data, we found that small and large clones showed differential expression in 132 genes at a false-discovery rate of 10% (**Fig. 4*B***). The variance among samples reflected the considerable heterogeneity of large clones within and between animals during microdissection (also shown in **Fig. 5*C***). Through gene set enrichment analysis, we found upregulation of multiple ciliated cell-related signatures in the small clones, indicating a highly differentiated status. We also detected an upregulation of the Ezh2 signature that is known for transcriptional repression (Cao et al., 2002; Vire et al., 2006) and down-regulation of stem-cell signature, further supporting a highly differentiated status of small clones (**Fig. 4*C***). In contrast, the large clones upregulated proliferation-related programs, including signatures for Mki67+ intestine cells and targets of B-Myb, which is a central regulator of cell proliferation (Musa et al., 2017) (**Fig. 4*D***). We did not detect canonical signatures of cell cycle activity, possibly because large clones contained only a small portion of slowly proliferating cells, as shown in our data below. Collectively, the sequencing data indicate that large clones are more proliferative and less differentiated compared to small clones. We also found that the large clones comprise both AcTUB+ ciliated cells and Pax8+ cells (**Fig. S5**), indicating the bi-potency of the large clone founder cells. In summary, these results support that founders of the large clones represent a more primitive FT subset.

**Figure 4:**
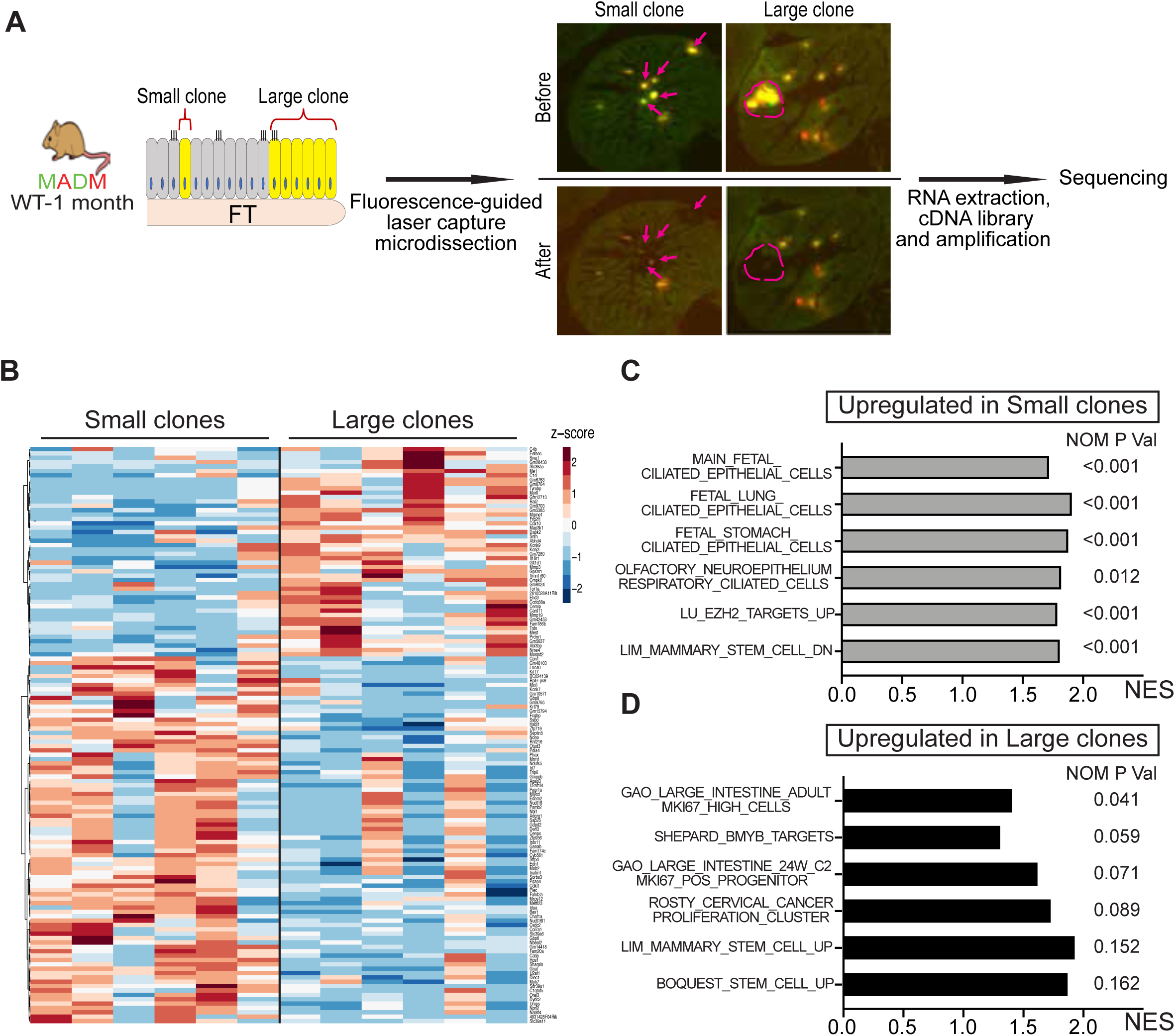
Founders of highly expansive clones represent a more primitive FT population. (A) The scheme of fluorescence-guided laser-capture microdissection of small and large clones from 1-month-old MADM-wildtype mice (*n*=6), and cDNA library preparation for sequencing. (B) Clustering of the 132 differentially expressed genes between small and large clones. log2 +1 transformed TMPs were subjected to Z-score normalization, and clustering was performed using Euclidian distance and the Ward.D method. (C) Gene set enrichment assay showing upregulation of gene sets related to ciliated cells in small clones, (D) Gene sets related to cell proliferation and stemness are upregulated in large clones.

**Figure 5:**
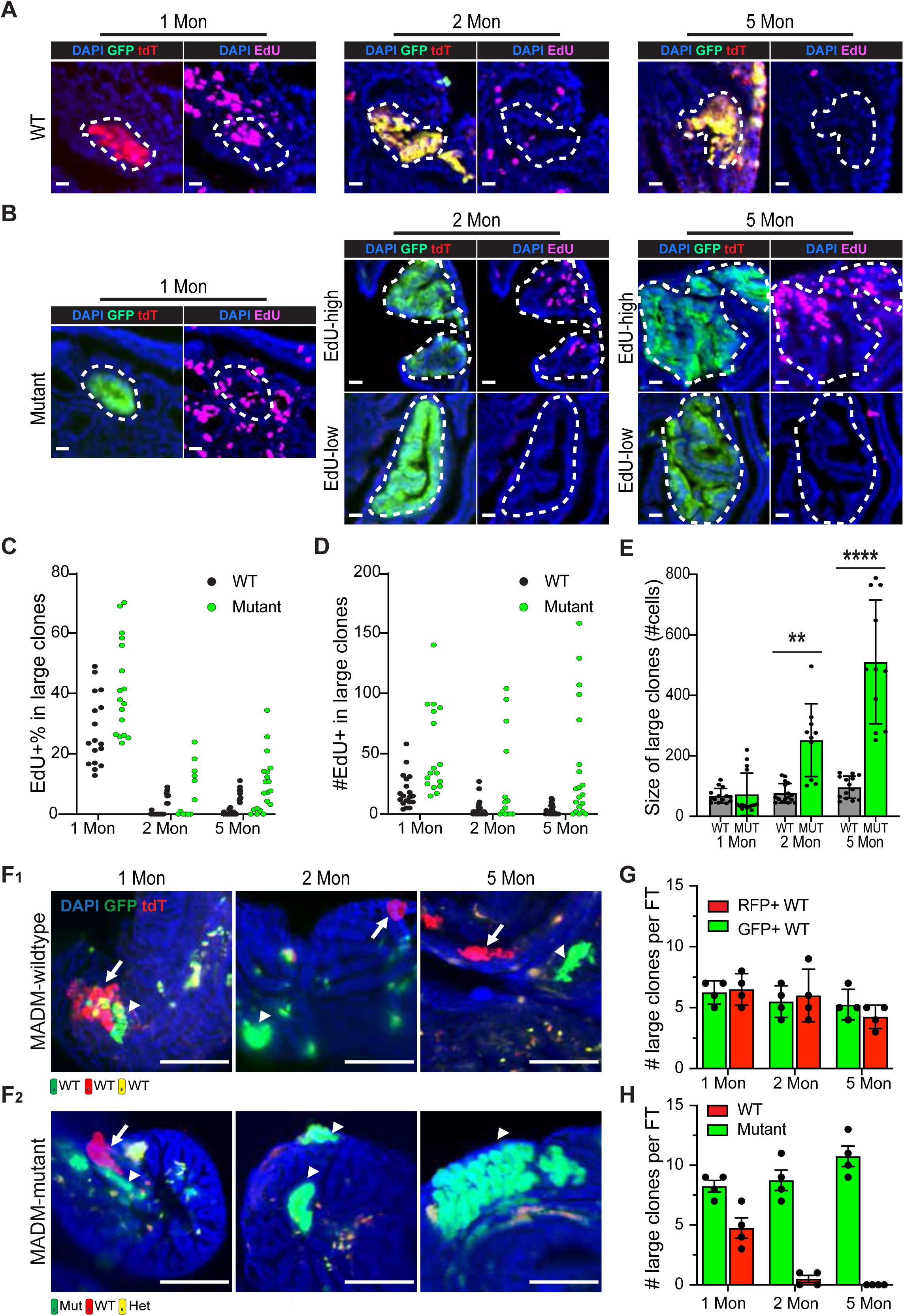
Oncogenic mutations prolonged expansion and persistence of large clones. (A) Representative images of EdU labeling in large wildtype clones at 1,2,5 months. Scale bar=20µm. (B) Representative images of EdU labeling in large mutant clones at 1,2,5 months. Scale bar=20µm. (C) EdU+% in each large wildtype and mutant clones at 1,2,5 month. Data pooled from three mice. (D) Number of EdU+ cells in each individual large wildtype and mutant clones at 1,2,5 month. Data pooled from three mice. (E) The average size of large wildtype clones and mutant clones from 1,2,5 months old mice. ± SEM. t-test, **<0.01****<0.0001. (F) F_1:_ Representative images of large GFP+ or RFP+ wildtype clones in 1,2,5 months-old MADM-wildtype mice. The arrow shows large RFP+ wildtype clones and the arrowhead shows large GFP+ wildtype clones. F_2:_ Representative images of large GFP+ mutant clones accompanied by large RFP+ wildtype clones in one-month-old MADM-mutant mice. Large RFP+ wildtype clones were lost at 2 & 5 months old. The arrow shows large RFP+ wildtype clones and the arrowhead shows large GFP+ mutant clones. Scale bar=50µm. (G) The number of large GFP+ and RFP+ wildtype clones per FT in MADM-wildtype mice at 1,2,5 months (n=4). (H) The number of large GFP+ mutant clones and large RFP+ wildtype clones per FT in MADM-mutant mice at 1,2,5 month (n=4).

### Oncogenic mutations prolonged the expansion and persistence of large clones

To understand how oncogenic mutations alter the clonal behavior of those FT Pax8+ cells with high clonogenic potential, we compared the proliferative activity of large clones with or without oncogenic mutations over time. To identify proliferating cells, we treated the mice with EdU in the drinking water for 7 days prior to harvest. The wildtype large clones showed a high proliferative rate at 1 month (EdU+:27.6±2.8%), which then dropped precipitously at 2 months (1.9±0.6%) and 5 months (2.0±0.5%). (**Fig. 5*A*, *C, D***), suggesting that the initial expansion capacity diminished quickly after the end of FT development (Ghosh et al., 2017). When oncogenic mutations were introduced, while the overall proliferative rate of large clones followed a similar diminishing trend after 1 month, we noted two important differences. First, the overall proliferation of mutant clones was maintained at a higher level than the wildtype ones at 2 and 5 months (**Fig. 5*B, C, D***). Second, the proliferative rate of large mutant clones at 2 and 5 months of age showed a large variance when assessed clonally. Some clones were nearly as active as 1-month and others almost completely stopped (**Fig. 5*B, C, D***, **Fig. S6**). These data indicate that oncogenic mutations could prolong proliferation in the large clones after the developmental window albeit not at full penetrance. Consequently, although mutant and wildtype large clones are comparable in size at 1 month, a portion of the mutant clones became much larger than wildtype ones at older ages (**Fig. 5*E***).

One of the unique strengths of genetic mosaic with MADM is that GFP+ mutant cells and RFP+ wildtype sibling cells are simultaneously generated in the same MADM-mutant animal (**Fig. 1*A_1_***), providing the opportunity to directly compare the clonal behavior of the mutant cells and wildtype cells in the same environment. Meanwhile, the MADM-mutant animals also have MADM-wildtype littermates (**Fig. S1*C***). Both types of animals contain wildtype clones, but one is surrounded by heterozygous/homozygous-mutant cells and the other is surrounded by wildtype cells, allowing us to dissect how mutant cells affect the clonal dynamics of wildtype cells. In MADM-wildtype animals, we found the large wildtype clones persisted long-termly (**Fig. 5*F_1_***). The number of GFP+ and RFP+ large clones are comparable at 1, 2, and 5 months. The total number of GFP+/RFP+ large clones also didn’t significantly change over time (**Fig. 5*G***). In MADM-mutant animals, although RFP+ large wildtype clones were present at one-month-old, they were seldom observed at 2 and 5 months old in more than 20 animals (**Fig. 5*F_2_, H***; **Fig. 2*C, F***). These data suggest that wildtype clones cannot persist in mutant FT, likely due to certain disadvantages in comparison to mutant clones.

Collectively, these results suggest that although the initial wave of clonal expansion is independent of oncogenic mutations, the mutations indeed prolong the proliferation of large clones and provide an advantage for their long-term persistence.

### Unbalanced differentiation parallels the further progression of mutant clones toward FT lesions

The epithelium of the normal distal FT consists of interspersed Pax8+ cells (~30%) and ciliated cells (~70%) (**Fig. S7*A, B***) (Harwalkar et al., 2021). On the contrary, precursor lesions in human FT such as “secretory-cell-outgrowths” (SCOUTs) and serous tubal intraepithelial carcinomas (STICs) predominantly consist of an uninterrupted stretch of Pax8+ cells (Chen et al., 2010; Ghosh et al., 2017). The loss of ciliated cells in FT is associated with a higher risk of developing ovarian cancer (Ghosh et al., 2017; Mehra et al., 2011; Tao et al., 2020). We asked whether the expanded mutant clones, as a candidate intermediate between normal and disease, could reflect this cell composition shift toward more Pax8+ cells. Indeed, we observed uninterrupted stretches of Pax8+ cells in expanded mutant clones reminiscent of early-stage abnormalities in humans, which contrasted with the interspersed Pax8+ and Pax8-cells in the adjacent non-mutant regions (**Fig. 6*A***). As a result, the proportion of Pax8+ cells in the progressive mutant clones was significantly higher than adjacent non-mutant regions (**Fig. 6*B***). We hypothesized that the uninterrupted stretches of Pax8+ cells result from a biased propensity of proliferating mutant cells to differentiate toward non-ciliated fate. To test this hypothesis, we performed an EdU pulse-chase experiment to determine the fate of newly born cells within progressive clones (**Fig. 6*C***). First, we mapped the length of EdU pulse, and found that 3-day administration led to clonal labeling of proliferating cells (**Fig. S7*C-G***). Then we quantified EdU+ clonal sizes after a 4-day chase and found that, while most wildtype clones remained at 1-2 cells, many mutant clones grew much larger (**Fig. S7*H-J***). When we examined the fate of cells in these EdU+ subclones, we found that, while 70% of cells in wildtype subclones had cilia, mutant EdU+ subclones were much less frequently ciliated (**Fig. 6*D, E***), suggesting a high propensity of mutant cells to adopt a non-ciliated cell fate. This altered cell differentiation fate may underscore early progression toward FT lesions.

**Figure 6:**
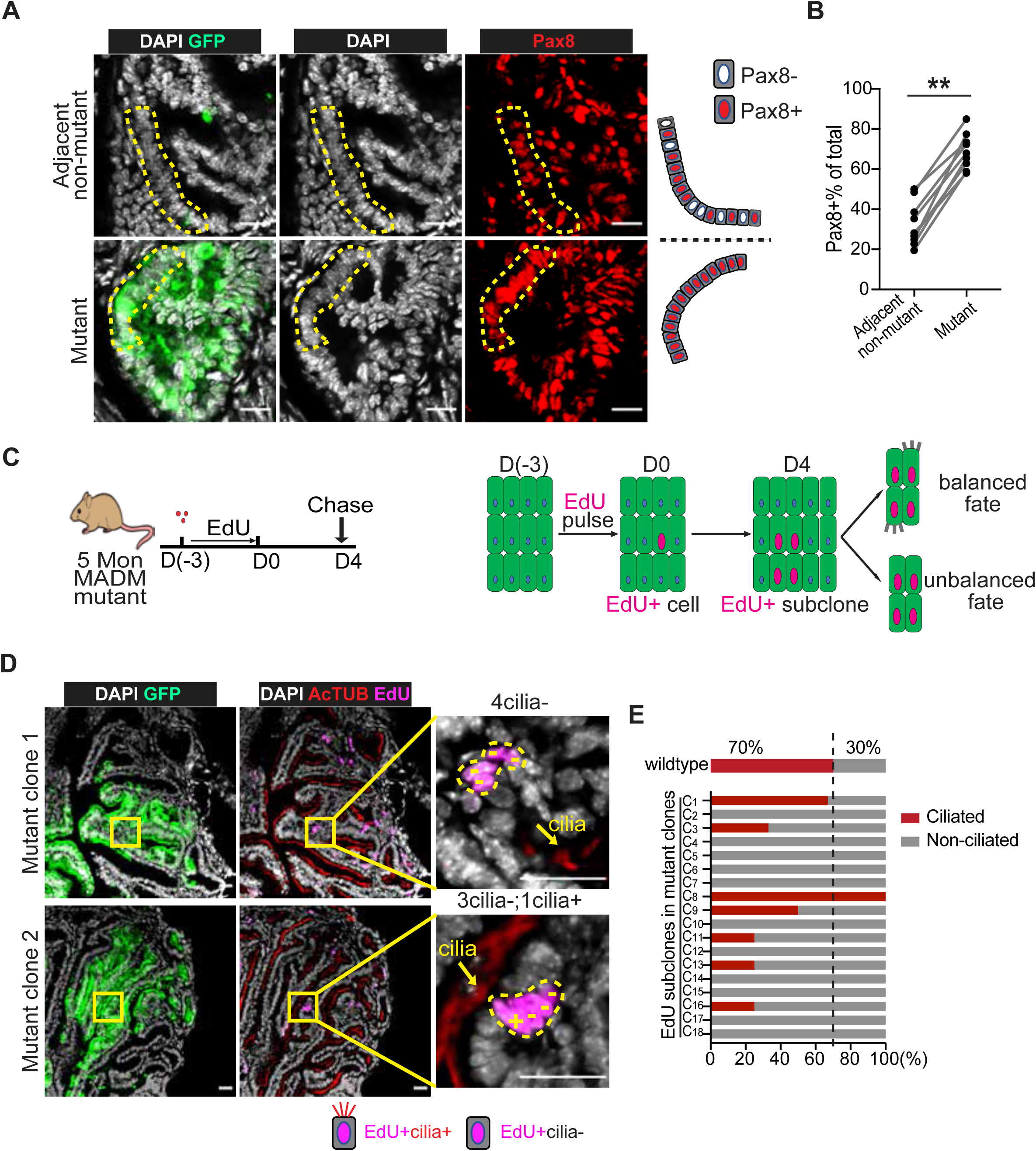
Unbalanced differentiation parallels the further progression of mutant clones toward FT lesions. (A) Representative images of Pax8+ cell distribution in progressive mutant clones (lower panel), and in adjacent non-mutant regions (upper panel). The mutant clones showed continuous expansion of Pax8+ cells. Scale bar=20µm. (B) The progressive mutant clones contained a higher portion of Pax8+ cells than adjacent non-mutant regions. Three mutant/non-mutant pairs were quantified from each of the three mice in total. Wilcoxon-test, **<0.01. (C) The scheme to perform clonal tracing of proliferating cells within progressive clones through EdU labeling, and potential outcomes. (D) Representative images show the cellular composition of EdU subclones within progressive mutant clones. The upper panel shows a four-cell subclone with only non-ciliated cells. The lower panel shows a mixture subclone with one ciliated cell (+) and three non-ciliated cells (-). Scale bar=25µm. (E) The cellular composition of EdU+ spots (>2 cells) within the mutant clones. In wildtype distal FT, ciliated cells account for the majority, however, in the EdU+ subclones within mutant clones, non-ciliated cells account for the majority.

## Discussion

It is extremely challenging to identify the cancer cell of origin in a tissue type with limited knowledge of lineage hierarchy. While FT Pax8+ cells can give rise to HGSOC, whether or not the Pax8 population consists of distinct cell types with unique tumorigenic potential remains unknown. In this study, we used a mouse genetic system called MADM to investigate the clonal progression of seemingly identical Pax8+ cells in the FT during the initiation stage of HGSOC. Despite limited prior knowledge, the dichotomous clonal expansion revealed by MADM clearly demonstrated that founder cells for progressive clones carry unique properties distinct from the majority of Pax8+ cells. This notion was further supported by the dichotomous clonal expansion of WT Pax8+ cells that are free of oncogenic mutations. Finally, we found that oncogenic mutations promoted cancer initiation of progressive clones by prolonging proliferation and shifting the differentiating trajectory. By dissecting cellular heterogeneity in cancer-initiating capacity, we found that HGSOC-relevant mutations can only promote cancer initiation in cells with permissive intrinsic properties.

### MADM enables clonal level analysis of the progression of individual mutant cells

Clonal analysis is an ideal approach to identify cancer cell-of-origin from a population of apparently similar cells. Although clonal level labeling can be achieved in mice containing both inducible Cre (CreER^T^ or tetracycline-inducible Cre) and Cre reporter transgenes by lowering the dosage of the Cre-inducing compound (Cheon and Orsulic, 2011), simultaneously knocking out tumor suppressor gene(s) in labeled cells is extremely difficult because recombination events that turn on Cre reporter are often uncoupled from the excision of the floxed gene(s), especially when Cre activity is tuned down to ensure clonality (Holzenberger et al., 2000; Muzumdar et al., 2007). Labeled cells could be wildtype; while mutant cells may not be labeled. In stark contrast, MADM achieves simultaneous gene knockout and reporter labeling in a single mitotic recombination event, ensuring a faithful coupling of mutant genotype and labeling. Additionally, the clonality is guaranteed without the need to tune down Cre activity because the probability of inter-chromosomal recombination is intrinsically low (Gao et al., 2014; Xu et al., 2014). Furthermore, the analytical power of MADM-based clonal tracking is greatly enhanced by the internal control, i.e., RFP-labeled wildtype sibling cells, which provide the opportunity to directly compare the clonal behavior of mutant cells with their wildtype siblings side by side, allowing one to detect even the subtlest abnormality in mutant clones (Beattie et al., 2017; Terry et al., 2020). Finally, cell status markers could be combined with MADM-labeling to reveal the status of proliferation, differentiation, and cell death of each cell within mutant clones, providing a snapshot of complex individual cell fate behind clonal progression (Driessens et al., 2012). When combined with spatial profiling (e.g. laser-capture microdissection followed by sequencing, or Nanostring GeoMx platform), one can dive deeply into the molecular details of individual clones (Yao et al., 2020). As a modular system, MADM-based clonal tracking of mutant cells at the cancer initiation stage can be applied to all tissues and cancer types, as a genome-wide library of MADM mice has recently been made available (Contreras et al., 2021).

### MADM reveals a rare, primitive Pax8+ subset enriched in the distal FT with cancer-initiating capacity

While FT Pax8+ cells have been identified as a cancer cell-of-origin for HGSOC (Lee et al., 2007; Perets et al., 2013; Zhang et al., 2019), we asked whether all or only a subset of the FT Pax8+ cells could initiate HGSOC. Quantitative clonal analysis of individual GFP-labeled mutant cells with MADM revealed a dichotomous size distribution: only a small proportion of Pax8+ cells are capable of clonal expansion after oncogenic mutations while others barely expand. These highly clonogenic cells are spatially enriched at the distal end of FT (close to the ovary), coinciding with the reported location of putative FT stem/progenitor-like cells based on organoid forming assays (Kessler et al., 2015; Paik et al., 2012; Xie et al., 2018). The fact that similar dichotomous clonal size in wildtype FTs free of mutations suggests that the founder cell for these clones could be stem/progenitor-like cells, considering that tissue stem cells often serve as cancer cell-of-origin (Greaves and Maley, 2012; Merlo et al., 2006; Visvader, 2011). This possibility is further supported by both the bi-potency of highly clonogenic cells based on histological analysis and the primitive status revealed by spatial profiling of the large clones. While cellular heterogeneity among FT Pax8+ cells has been revealed by both our spatial profiling data and recent single cell sequencing data (Dinh et al., 2021; Hu et al., 2020; Johnson et al., 2022; Ulrich et al., 2022), there are very limited overlaps in the putative stem/progenitor-like signatures from these studies, likely due to exceptional heterogeneity among the Pax8 population, technical limitations, or both. For example, our spatial profiling experiment is most likely confounded by the fact that most cells within the large clones have differentiated into secretory or ciliated cells, thus diluting the stem signatures of the clonal founders. In the future, spatial profiling of large clones at the single-cell resolution should provide a much better chance of identifying markers for the highly clonogenic stem/progenitor-like cells.

### Oncogenic mutations drive clonal progression in a permissive context

Both oncogenic mutations and the lineage programs inherent in the cells where mutations occur are critical for cancer initiation (Greaves and Maley, 2012; Lytle et al., 2018; Visvader, 2011). Our work suggests that HGSOC-relevant mutations cannot induce clonal progression in a non-permissive context since the small clones never took off. However, in a permissive context, mutations powerfully drive clonal progression by lengthening the duration of cell proliferation, as indicated by clonal sizes and proliferation rates between MADM-WT and MADM-mutant FTs along a time course. Notably, the proliferation-promoting effect of mutations in large clones varies greatly, implying an inherent tendency for even the permissive clones to halt their progression. Besides cell proliferation, mutant cells in expanded clones also manifested biased differentiation toward a Pax8+ fate, resulting in stretches/nodules of uninterrupted Pax8 cells and a significantly decreased proportion of ciliated cells, a feature reminiscent of human precursor lesions known as SCOUTs and STICs (Chen et al., 2010; Ghosh et al., 2017; Tao et al., 2020). Because the Pax8+ population contains FT stem/progenitor-like cells based on previous lineage tracing studies (Ghosh et al., 2017), this biased cell fate may reflect a tendency of oncogenic mutations to preserve stemness, a feature often found during the progression of many cancer types (Alcolea et al., 2014). These observations warrant further investigations into the baseline signaling network in FT stem cells as well as the interactions between this network and oncogenic mutations that lead to augmented proliferation and biased differentiation.

### Implications of premalignant heterogeneity for cancer early-detection and prevention studies

Although the exact differences between the highly expansive and stalled Pax8+ cells need to be further explored, the clonal dynamics revealed by our study carry important implications for early detection and prevention studies of HGSOC. We uncovered a gradual attrition process of mutant Pax8+ cells during cancer initiation and premalignant progression (**Fig. S8**). Initially, the vast majority of mutant cells stalled while only a small portion enriched at the distal end of FT expanded. As time goes by, among initially expanded clones, only some sustained their progressiveness. Moreover, within continuously progressing mutant clones, only a small fraction of the cells maintained proliferative activity. This discovery calls for careful consideration for cancer early detection studies: one must hone in the rare proliferating cells within continuously progressing mutant clones to discover biomarkers to guard against the risk of over-diagnosis. Indiscriminative profiling of all mutant cells would bring in tremendous noise from stalled mutant cells and mask the true signal from cells capable of further progression to malignancy (Allred et al., 2008; Greaves and Maley, 2012; Park et al., 2010). For the same reason, to measure the efficacy of potential cancer prevention strategies, one should home in on progressing cells rather than the bulk of mutant cells of which most have already reached their dead end.

In summary, our study demonstrates a new application of a mouse genetic mosaic system to dissect cellular heterogeneity of cancer-initiating capacity in tissues with limited knowledge of lineage hierarchy. For HGSOC, the work reveals cellular alterations underlying disease progression at an unprecedentedly early stage. Broader implementation of this system should be valuable in deepening our understanding of cancer initiation, paving the way for effective cancer early detection and prevention in many cancer types.

## Acknowledgments

We thank Dr. Zhe Li, Xiaoyu Zhao, and Bing Xu for providing critical feedback on the manuscript and Jocelyn Ray for the artwork of the fallopian tube cartoon. We also thank Dr. Stacey Criswell at the Advanced Microscopy Facility, Dr. Pat Pramoonjago at the Biorepository and Tissue Research Facility, and Sheri Vanhoose at the Research Histology Core, and Shelly Verling at the vivarium for their assistance on the project. These core facilities are supported by UVA Cancer Center grant #P30-CA044579. We are grateful to Dr. Ammasi Periasamy and Dr. Ruofan Cao at the Keck Center for the usage of the Light-sheet Z.1 microscopy system. This work was partly supported by the Department of Defense Ovarian Cancer Research Program #W81XWH-17-1-0174 (J.K.S-D. & H.Z.), the Rivkin Center (H.Z.), the National Cancer Institute #R01-CA256199 (K.A.J. & H.Z.) and #R50-CA265089 (L.W.), the UVA Cancer Center Seed Grant (J.K.S-D. & H.Z.), the UVA Systems & Biomolecular Data Sciences Training Grant (A.C.AY), and the UVA Cancer Center Training Grant (J.Z.).

## Author Contribution

Conceptualization, J.Z., J.K.SD., and H.Z.; Methodology: J.Z., E.C., Y.J., L.W., and B.E.K.; Data Collection & Analysis: J.Z., A.C.AY., E.K., T.J., K.A.A, K.A.J, and H.Z.; Writing-Review & Editing: J.Z., A.C.AY., K.A.J, J.K.SD., and H.Z.; Funding Acquisition: J.K.SD., and H.Z.

## Figure Legends

**Figure S1:**
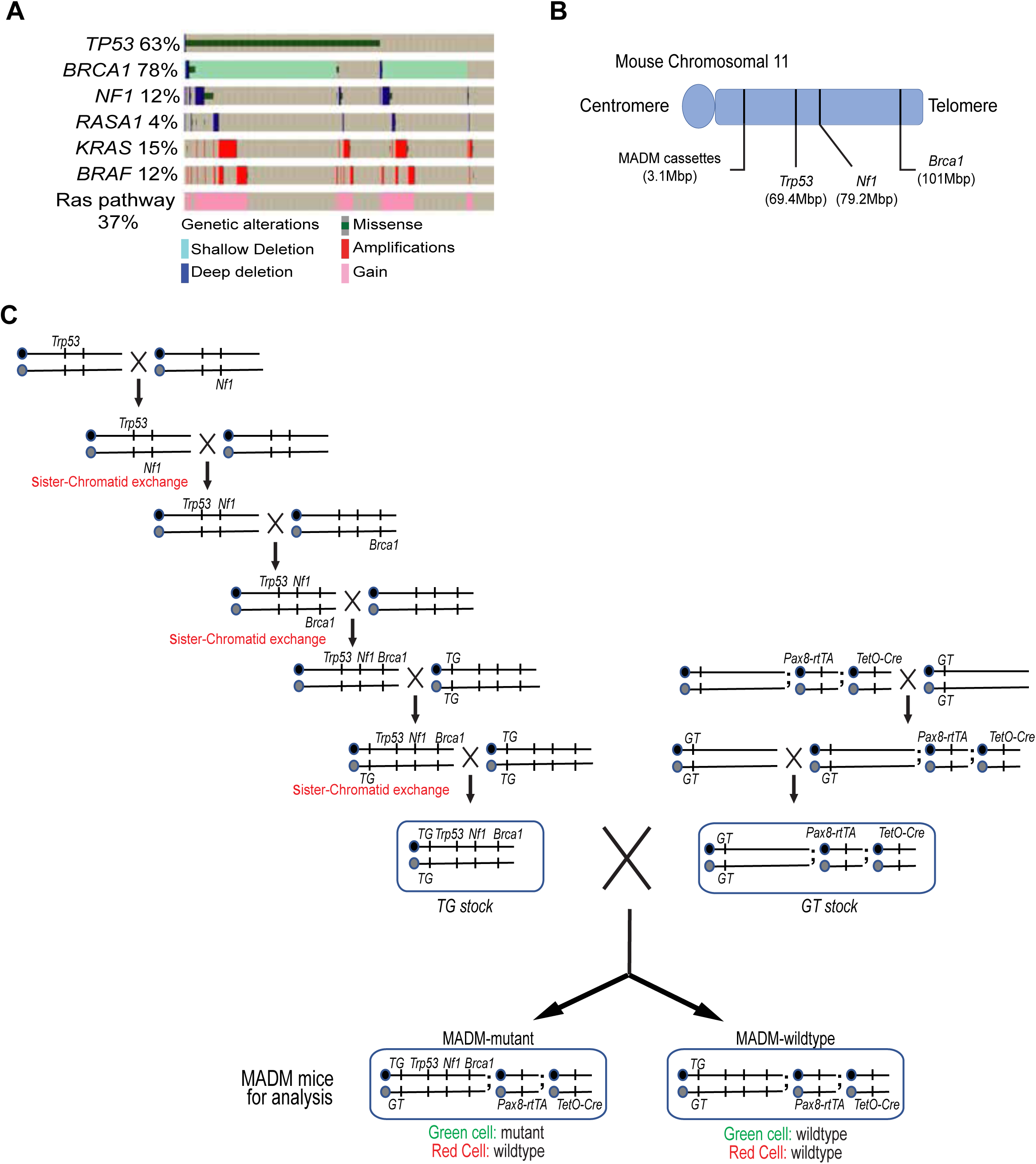
Additional details for establishing a MADM-based mouse model with scattered GFP+ mutant cells in the FT. (A) Assessment of HGSOC mutation spectrum with TCGA datasets. (B) Chromosomal locations of *MADM cassettes*, *Trp53, Nf1,* and *Brca1* on mouse chromosome 11. The physical locations were indicated. (C) The scheme to breed MADM-mutant mice and MADM-wildtype mice. We maintain two separate stocks and produce MADM mice by intercrossing the stocks. *Trp53, Nf1,* and *Brca1* mutations were incorporated into The TG stock, and the Cre transgenes (*Pax8-rtTA, TetO-Cre*) were incorporated into the GT stock.

**Figure S2:**
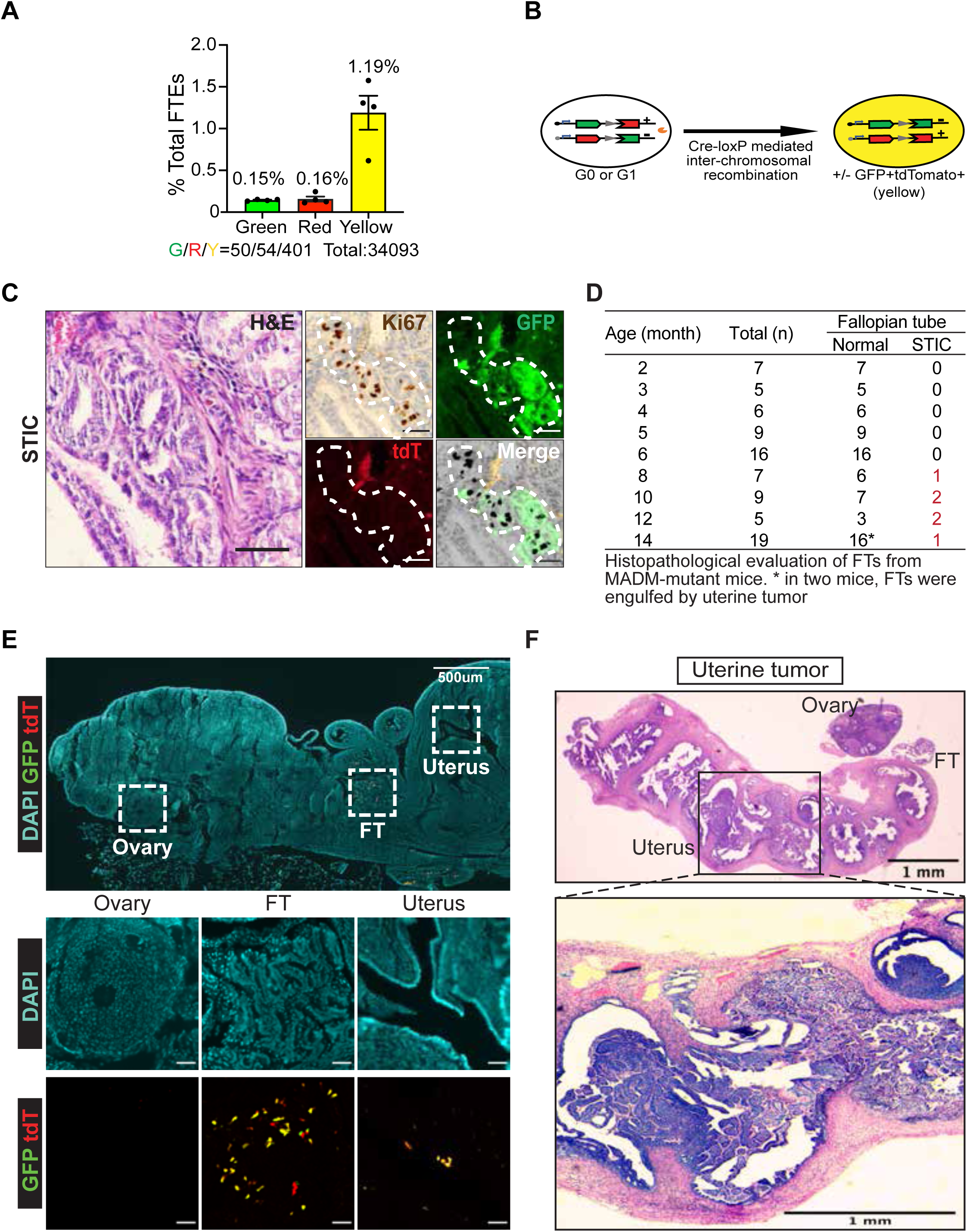
Formation of FT STICs lesions and uterine tumors in the MADM model. (A) Frequency of MADM-labeled cells in FT after inducing Cre between postnatal day 0 and day 21 (P0-21) (±SEM, *n*=3). (B) The percentage of yellow cells is much higher because Cre-mediated inter-chromosomal recombination occurring in G1 or post-mitotic cells (G0) generates yellow cells without altering genotype. (C) H&E staining and Immunofluorescence staining show FT STICs that overlap with GFP, but not tdTomato. These GFP+ lesions are composed of foci of atypical columnar epithelial cells lining the papillary structures of the fallopian and demonstrate enlarged, crowded nuclei with hyperchromasia, prominent nucleoli, scant cytoplasm, and a lack of cilia. Immunohistochemistry staining for Ki-67 demonstrated an increase in proliferative activity as compared to the normal adjacent epithelium. Scale bar=50µm. (D) STICs frequency in the FTs from MADM-mutant mice of an age cohort. (E) MADM-labeled cells were found in both the FT and the uterus, but not in the ovary. (F) Formation of uterine tumors at around 12 months.

**Figure S3:**
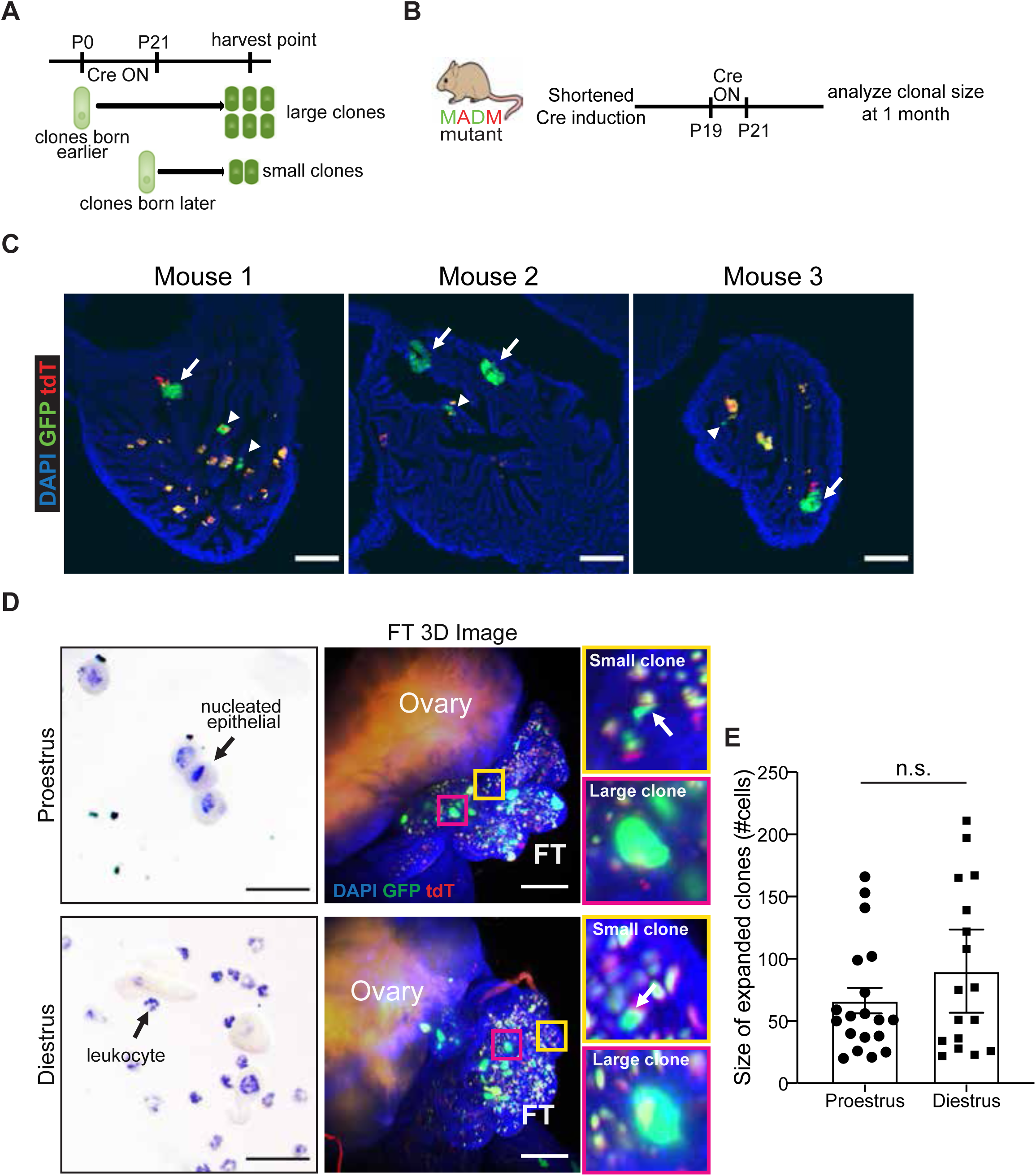
Clonal size heterogeneity is not simply due to unsynchronized clonal age, and is not dramatically affected by the estrus cycle. (A) The clonal age hypothesis: clones born earlier within the P0-21 range will show larger size at the harvest point, whereas clones born later will show smaller size. (B) The scheme of Dox treatment to synchronize the clonal age. (C) Representative images show the heterogeneous size of MADM-labeled clones in age-synchronized mice (*n*=3). The arrow shows expanded clones and the arrowhead shows non-expanded clones. Scale bar=100µm. (D) FTs were collected at proestrus or diestrus in post-puberty MADM-mutant mice (~10 weeks, *n*=3 each), representative 3D images of whole FT show co-existence of non-expanded GFP+ mutant clones (yellow box) and expanded GFP+ mutant clones (magenta box). Scale bar=100µm. (E) The clonal size of expanded mutant clones from mice harvested at proestrus and diestrus. ±SEM. t-test, p=0.21.

**Figure S4:**
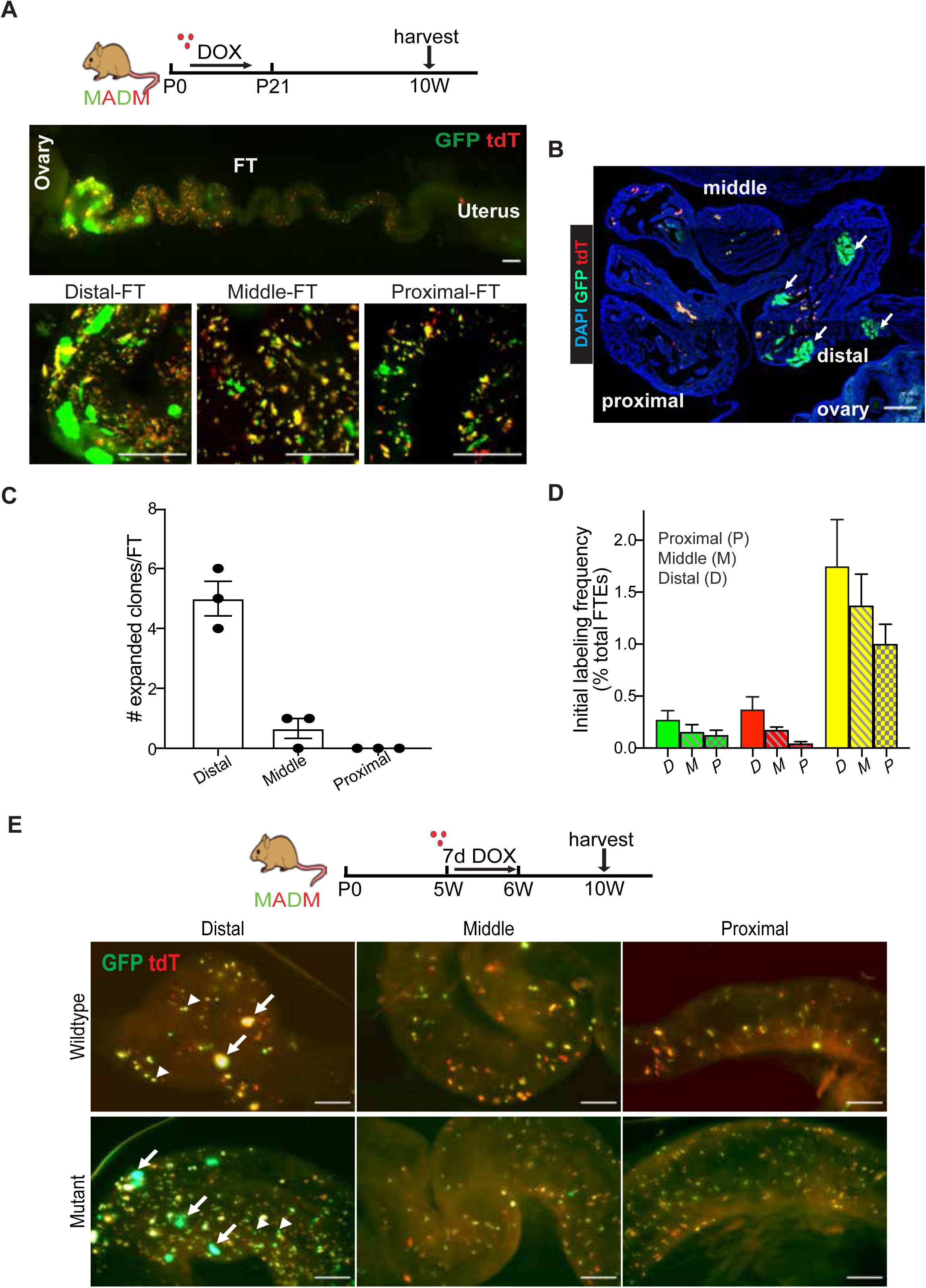
Spatial distribution of expanded clones in MADM-mutant mice; MADM-labeling frequency in each FT segment initially; and spatial distribution of expanded clones when induced post-developmentally. (A) Images of whole-mount FT images from 10 weeks old mutant FTs (*n*=3), showing that expanded mutant clones were mainly in the distal region. Scale bar=200µm. (B) Sectioned FT from 10 weeks old mutant FTs showing a distal preference for expanded mutant clones. Scale bar=100µm. (C) Quantification of the number of expanded clones in distal(D), middle(M), and proximal(P) regions from 10 weeks old MADM-mutant mice. (±SEM, *n*=3). (D) Initial MADM labeling frequency in distal(D), middle(M), and proximal(P) FT regions right after P0-21 Doxycycline administration (*n*=3). (E) Inducing MADM labeling in fully differentiated FTs: Doxycycline was administered to MADM-wildtype/mutant mice between 5-6 weeks, then FTs were assessed at 10 weeks. Upper panel: wildtype FT; Lower panel: mutant FT. The arrow shows expanded large clones and the arrowhead shows non-expanded clones. Scale bar=200µm.

**Figure S5:**
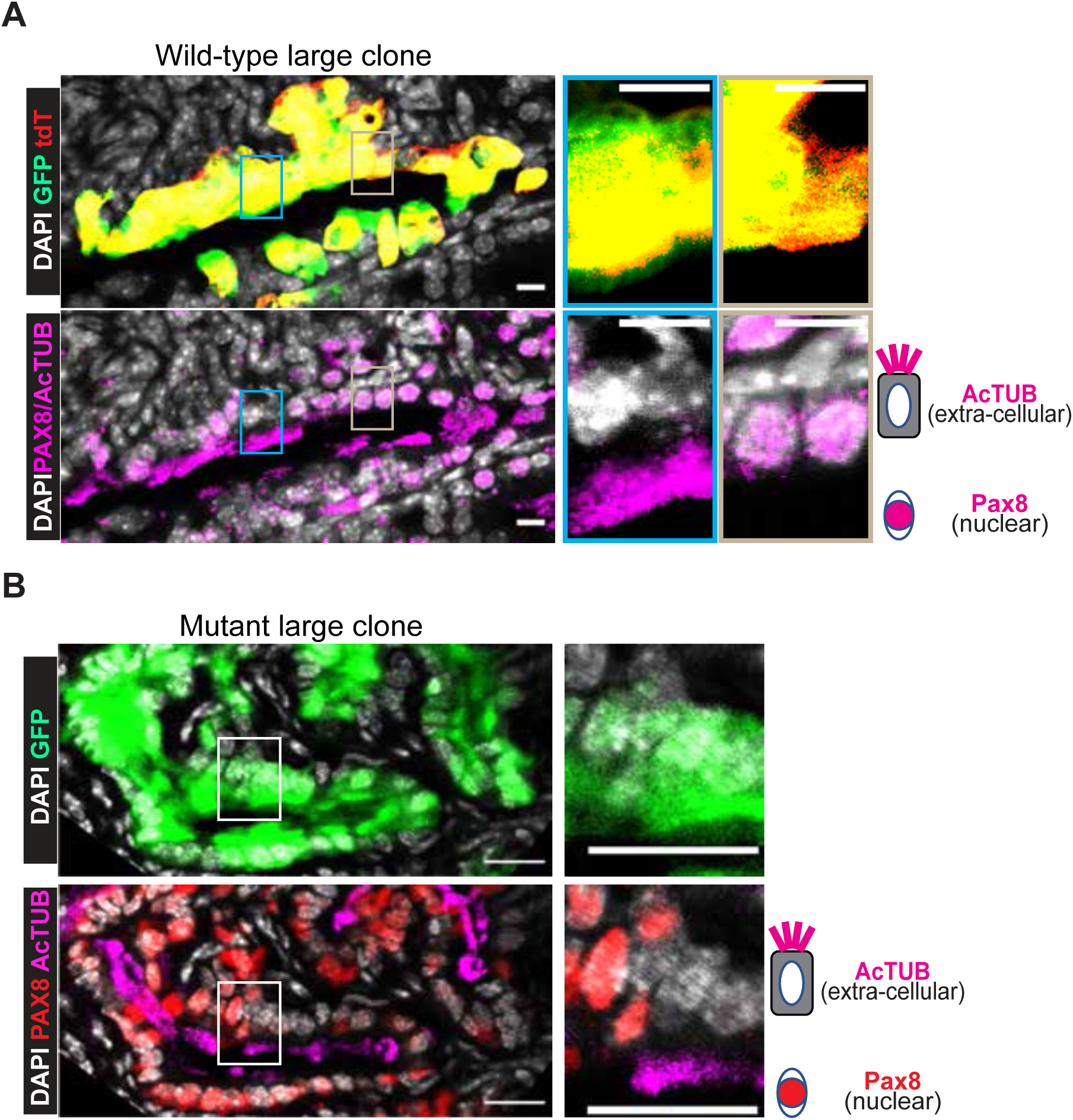
Expanded large clones consist of both AcTUB+ ciliated cells and Pax8+ cells. (A) Representative image of expanded wildtype clones that comprises both AcTUB+ ciliated cells and Pax8+ secretory cells. Scale bar=20µm. (B) Representative image of expanded mutant clones that comprises both AcTUB+ ciliated cells and Pax8+ secretory cells. Scale bar=20µm.

**Figure S6:**
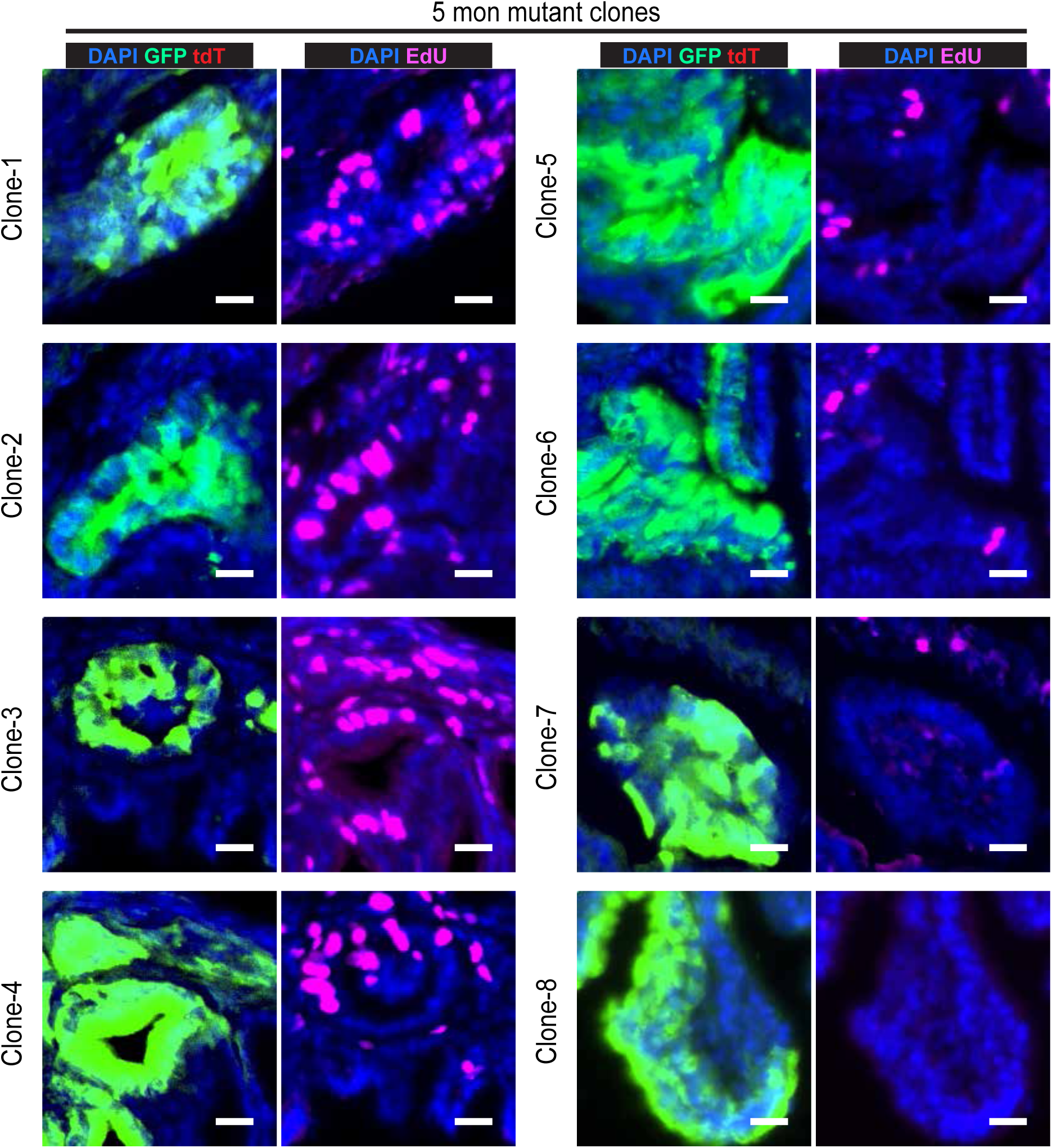
More images of EdU labeling in expanded mutant clones at 5 months. Scale bar=20µm.

**Figure S7:**
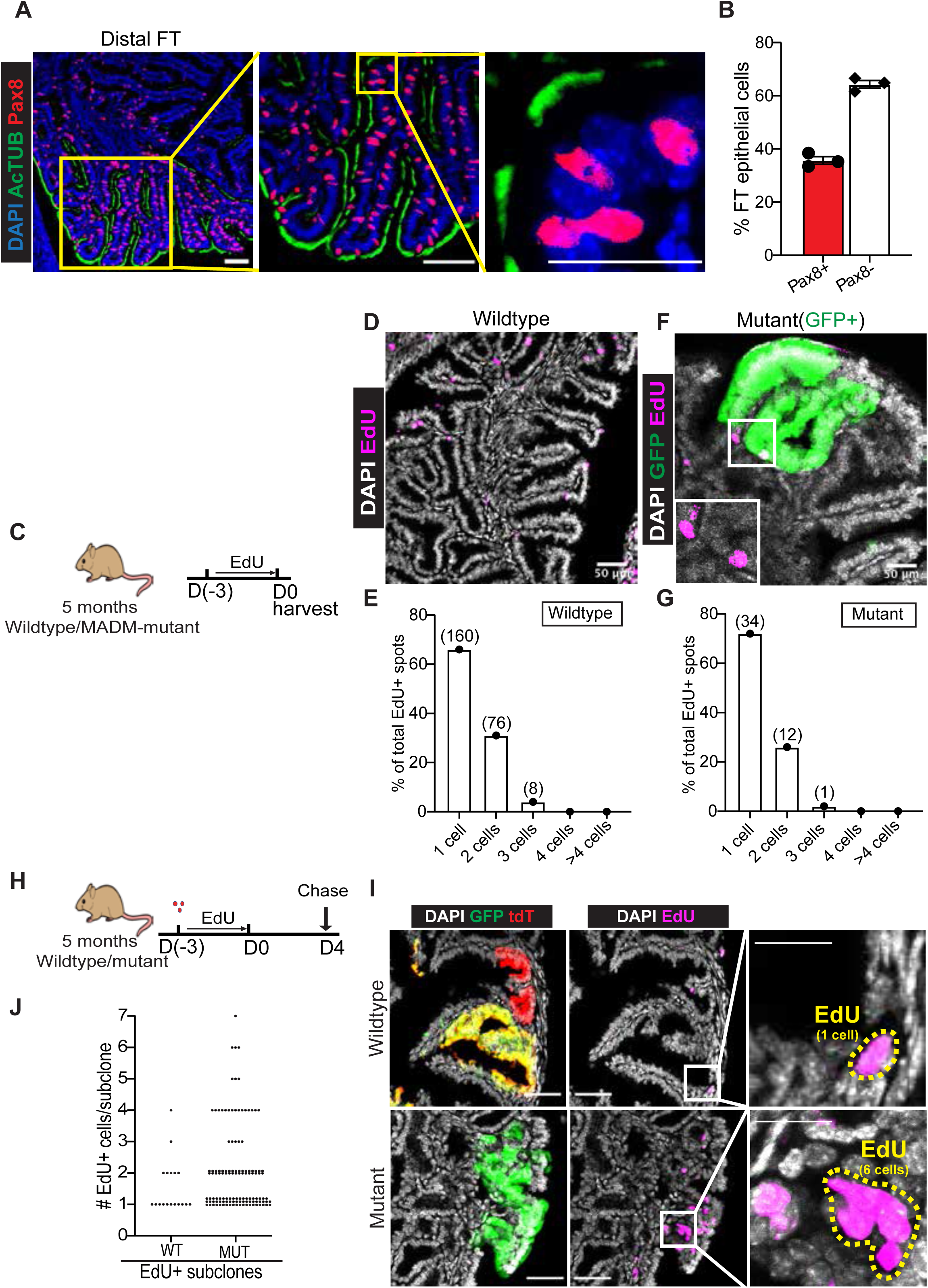
The Pax8+ cell distribution pattern in wildtype distal FT and 3-day EdU labeling marked cells at clonal density. (A) Alternating Pax8+ and AcTUB+ cells in the distal FT from 5 months wildtype mice. Scale bar=50µm for the left and middle panels, and scale bar=25 µm for the right panel. (B) Percentage of Pax8+ and Pax8-cells in the distal FT. (±SEM, *n*=3). (C) Scheme for EdU treatment and analysis for labeling proliferating cells at clonal density. (D) EdU staining in wildtype FTs revealed that EdU-labeled cells were scattered, and most EdU+ spots were shown as one-cell or two-cell status. Scale bar=50µm. Representative image from 3 mice. (E) The size distribution of EdU+ spots in FTs from 5 months old wildtype mice (*n*=3). The total number of EdU spots (n) is shown above the bar graph. (F) EdU staining in mutant FTs revealed that EdU-labeled cells were also scattered, and most EdU+ spots were still shown as one cell or two cells. Scale bar=50µm. Representative image from 3 mice. (G) The size distribution of all EdU+ spots in FTs from 5 months MADM-mutant mice (*n*=3). The total number of EdU+ spots (n) is shown above the bar graph. (H) Scheme for EdU pulse-chase assay. (I) Upper panel: an EdU+ clone of one cell within expanded wildtype clones when chased at day 4. Lower panel: an EdU+ clone of six cells within progressive mutant clones. Scale bar=20µm. (J) The size of each EdU+ clone within expanded wildtype clones and progressive mutant clones. Data pooled from three mice for each group.

**Figure S8:**
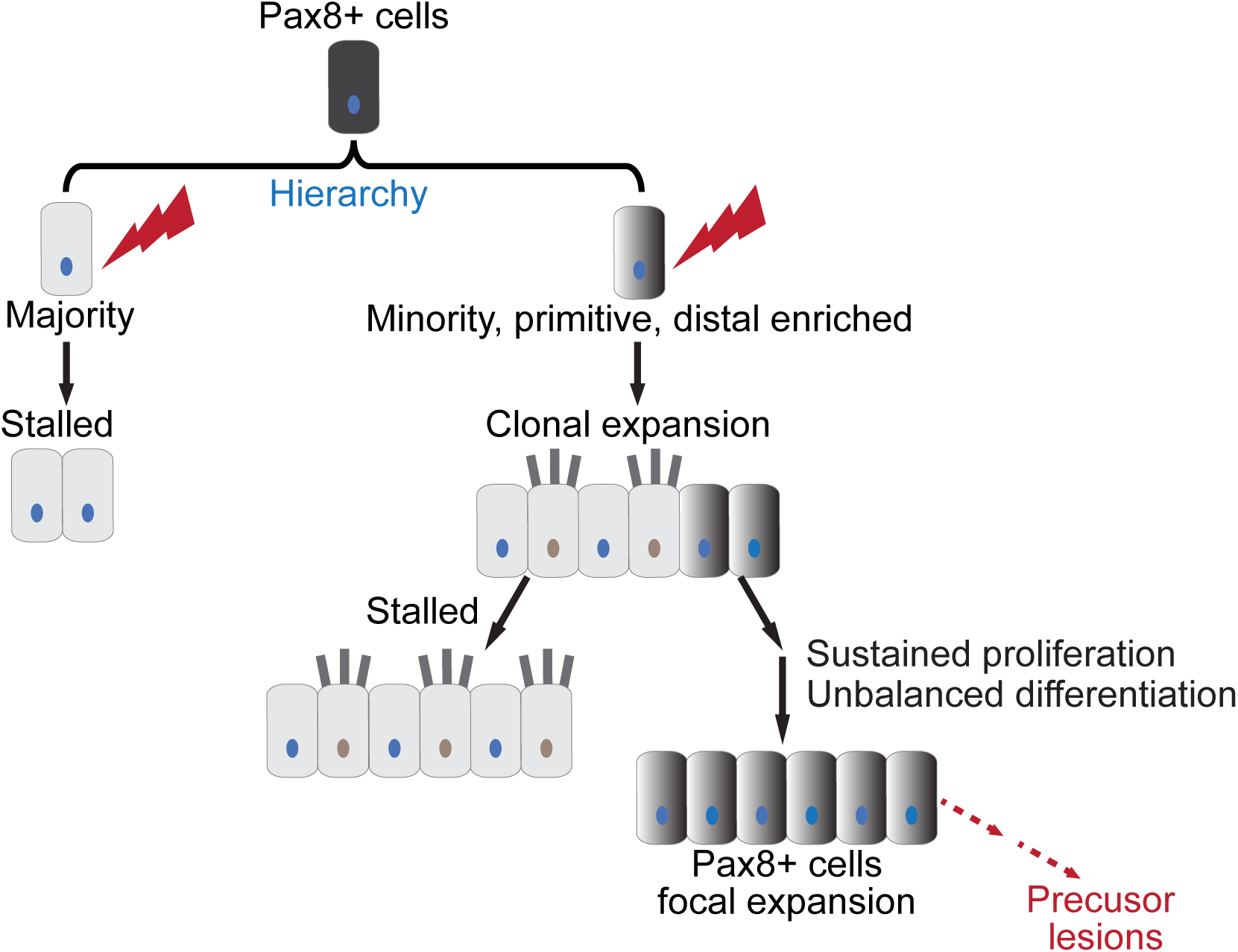
Model delineating heterogeneous ovarian cancer-initiating capacity of FT Pax8+ cells and their progressive fate: The initiated FT Pax8+ cells adopt distinct clonal expansion fate shortly after being generated, with the majority stalled, a minority which is spatially enriched in the distal FT can quickly expand. The initially expanded mutant clones then experience a further divergence in their clonal fate, either lose their progression or keep a long-term progression. The latter presents unbalanced differentiation and causes focal expansion of Pax8+ cells, which underlies further progression toward FT lesions.

## Star Methods

### Animals

All animal procedures are approved by the Institutional Animal Care and Use Committee (IACUC) at the University of Virginia in accordance with national guidelines to ensure the humanity of all animal experiments. The following mouse lines were crossed to establish the MADM-mutant and MADM-wildtype mice: *TG11ML* (stock NO. 022976 JAX), *GT11ML* (stock NO. 022977 JAX), *BRCA1^flox^* (strain NO. 01XC9 NCI), *NF1^flox^* (strain no. 01XM4; NCI), *Trp53^KO^*(stock no. 002101; JAX), *Pax8-rtTA* (stock NO. 007176 JAX)(Traykova-Brauch et al., 2008), *TetO-Cre* (stock NO. 006234 JAX) (Perl et al., 2002).

### In vivo drug delivery

Doxycycline was administered through the drinking water (2mg/ml) to the nursing mother, which transmits to MADM pups through milk (P0-P21) to induce Cre recombinase. 5-ethynyl-2’– deoxyuridine (Invitrogen, Cat# A10044) was administered through drinking water (0.5mg/ml). EdU staining was performed following standard procedures (Salic and Mitchison, 2008).

### Immunostaining

Reproductive tracts of MADM mice were harvested and then fixed overnight with cold 4% paraformaldehyde (PFA) at 4°C. Tissues were then washed with PBS to remove recessive PFA, soaked with 30% sucrose, embedded in optimal cutting temperature (OCT), and stored at −80°C. The tissues were sectioned at 18μm thickness. For staining, slides were incubated in Permeabilization/blocking buffer (0.3% Triton-X 100 in PBS [PBT] plus 5% normal donkey serum) for 20 min then primary antibodies (Pax8,1:50,10336-1-AP, Proteintech; AcTUB,1:500, T7451, Sigma) diluted in Permeabilization/block buffer were added and incubated at 4°C overnight. Secondary antibody incubation was performed for 1 hour at RT in PBT. To stain nuclei, slides were incubated in DAPI solution (1ug/mL in PBT) for 5 min before mounting with 70% glycerol and covered. Fluorescent images were acquired on Zeiss LSM 700/710 confocal microscope. Images were processed with Zen and Fiji.

### Tissue clearing with the CUBIC method and 3D imaging

The PFA fixed ovary and fallopian tube were cleared for large-scale 3D imaging with the standard CUBIC method (Susaki et al., 2015). Tissues were first immersed in 50% reagent-1 and shaken at 110 rpm, 37°C for 6h and then transferred to 100% reagent-1 with DAPI (1ug/ml) to shake for 2-3 days until becoming transparent. After reagent-1, tissues were washed three times with PBS, 30 mins each with shaking to remove the reagent-1. Tissues were immersed in 50% reagent-2 for 6h with shaking at 37°C; thereafter, the buffer was exchanged for 100% reagent-2 with shaking for 24h. The Zeiss Z.1 light-sheet microscopy system was used to acquire images. Tissues were embedded in 2% (w/w) agarose (LE quick dissolve agarose, GeneMate E-3119-500) gel; the agarose was dissolved in a modified reagent-2 (10% urea, 50% sucrose, 30% H_2_O, hot stir at 80°C until fully dissolved, then add 10% Tri-ethanolamine). The warm agarose gel solution together with tissue was aspirated into a 1ml syringe with the neck cut off, and then the syringe was placed on ice for quick solidification. The syringe was placed on the holder of the light-sheet microscope, and the tissue was pushed out for imaging.

### Counting clone size

<u>The whole</u> FT was cleared and imaged at 5 um intervals with light-sheet microscopy at single-cell resolution through the entire tissue. Three-dimensional reconstruction of clones was performed; labeled cells that were found at the exact location through multiple image stacks were counted as one clone. The size of clones was measured manually by counting the DAPI-stained nuclei.

### Mathematical detection of outlier clones

The clonal size distribution along the fallopian tube was evaluated in terms of cell division events. With n as the number of cells per clone, there will be n-1 cell divisions. For clonal size distribution at five months, we fitted with a negative binomial distribution, which captures the behavior of the over-dispersed data observed in our mouse models, by allowing the mean and variance to be different. Clonal size distribution at 1 and 2 months, which showed a smaller range, were fitted with a Poisson distribution. To estimate the statistical significance of our fitting, we generated null distributions for the Kolmogorov-Smirnov (KS) statistic between an ideal probability mass function and a random negative binomial/Poisson distribution over 1.000 iterations. The parameters used for generating the null distributions were obtained by merging all datasets and taking a maximum of 44/8-9-10, which showed significance in our first general screening. Then, the empirical nominal p-value of the observed KS for the merged data was calculated relative to the null distribution until a cutoff with significance was observed. Our final cutoff was set as the maximum number of divisions for which the probability of observing our KS was p >= 0.1. The validity of this cutoff was confirmed by evaluating the individual datasets over the null distributions. Then, the maximum expected clonal sizes (n) in the fallopian tube were calculated, and any clone with a size larger than n was taken as the outlier. Subsequently, we computed the expected clone sizes for all the models, and we tested their significance against the observed values in our datasets using Fisher exact/Chi-square tests. Finally, we defined over-proliferative clones (Expected-observed outliers) for the size distributions observed under the Poisson model and tested their significance between 1, 2, and 5 months by an Anova with Tukey’s Honest Significant Difference test.

### Fluorescence-guided laser-capture micro-dissection of fallopian tube epithelial cells

Cryo-embedded fresh FTs were sectioned at 8 µm thickness and were then dehydrated with 70% ethanol (30 sec), 95% ethanol (30 sec), and 100% ethanol for 1 min, followed by clearing with xylene (2 min). After air drying, slides were micro-dissected with Arcturus XT LCM instrument (Applied Biosystems) and Capsule HS caps (Arcturus). The typical instrument settings of ~50 mW power and ~2 ms duration with the smallest spot size were used.

### RNA extraction, reverse transcription, and amplification for quantitative PCR

The procedures were described before (Singh et al., 2019). In brief, RNA from the micro-dissected cells was extracted through enzymatic digestion with proteinase K (Sigma), then reverse transcribed with 5’-biotin modified oligo(dT)24 (IDT) and SuperScript III (50 °C, 30 min; then heat inactivation at 70 °C for 15 min). The first-strand products were purified with Streptavidin magnetic beads (Pierce) and Super Magnet Plate (Alpaqua) to remove genomic DNA. Then the cDNA was poly(A)-tailed and amplified with 25 cycles of PCR with AL1 primer: ATTGGATCCAGGCCGCTCTGGACAAAATATGAATTCTTTTTTTTTTTTTTTTTTTTTTTT

### RNA sequencing and analysis

For sequencing, the cDNA samples were re-amplified following protocols described before (Singh et al., 2019). 100 µl PCR reactions were set up for each sample: 1) 5 µg AL1 primer, 2) 1x High-Fidelity buffer (Roche), 3.5 mM MgCl2, 200 µM dNTPs, 100 µg/ml BSA, 3) 1 µl cDNA template and 4) 1 µl Expand High Fidelity polymerase (Sigma). The following PCR program was used: 1 min at 94 °C, 2 min at 42 °C, and 3 min at 72 °C. 10 cycles were run to keep an exponential phase of re-amplification (Wang and Janes, 2013). The products were purified with AMPure XP beads (Beckman Coulter) according to the manufacturer’s protocol to remove excessive primers. The concentration of purified cDNA libraries was measured with the Qubit dsDNA BR Assay Kit (Thermo Fisher). Libraries were diluted to 0.2 ng/µl for tagmentation with the NXTR XT DNA SMP Prep Kit (Illumina). Libraries were multiplexed at an equimolar ratio, and 1.3 pM of the multiplexed pool was sequenced on a NextSeq 500 instrument with a NextSeq 500/550 Mid/High Output v2.5 kit (Illumina) to obtain 75-bp paired-end reads and an average of 3.6 million alignments per sample. From the sequencing reads, adapters were trimmed using fastq-mcf in the EAutils package (version ea-utils.1.04.636), and with the following options: -q 10 -t 0.01 -k 0 (quality threshold 10, 0.01% occurrence frequency, no nucleotide skew causing cycle removal). Datasets were aligned to the mouse transcriptome (GRCm38.95), using RSEM (version 1.3.0) with the following options: --bowtie2 --single-cell-prior --paired-end (Bowtie2 transcriptome aligner, single-cell prior to account for dropouts, paired-end reads). RSEM read counts were converted to transcripts per million (TPM) by dividing each value by the total read count for each sample and multiplying by 10^6^. DESeq2 (version 1.37.0) was used to identify differentially expressed transcripts between the small and large clone groups. Genes with an Padj < 0.1 were considered significantly up- or downregulated between the two groups. To generate a heatmap for the differentially expressed genes between small and large clones, their TPMs were transformed with log2 +1 and then were subjected to Z-score normalization. Clustering was performed using Euclidian distance and the Ward.D method. Gene-Set Enrichment Analysis (GSEA) was implemented to determine molecular signatures for the differential expressed transcripts (“h.all.v7.4.symbols.gmt”). Sequencing data were deposited to the GEO database: GSE210409.

